# The impact of chemical pollution across major life transitions: a meta-analysis on oxidative stress in amphibians

**DOI:** 10.1101/2023.09.04.556172

**Authors:** Colette Martin, Pablo Capilla-Lasheras, Pat Monaghan, Pablo Burraco

**Affiliations:** School of Biodiversity, One Health and Veterinary Medicine, University of Glasgow, Glasgow G12 8QQ, United Kingdom; Doñana Biological Station (CSIC), 41092, Seville, Spain

**Keywords:** contamination, global change, life history, metamorphosis, oxidative damage, pollutants, trade-offs

## Abstract

Among human actions threatening biodiversity, the release of anthropogenic chemical pollutants -which have become ubiquitous in the environment- is a major concern. Chemical pollution can induce oxidative stress and damage by causing the overproduction of reactive oxygen species (ROS), and affecting the antioxidant system. In species undergoing metamorphosis (∼80% of all extant animal species), antioxidant responses to chemical pollution may differ between pre- and post-metamorphic stages. Here, we meta-analysed (N = 86 studies, k = 2,012 estimates) the impact of chemical pollution on the redox balance across the three major amphibian life stages (embryo, larva, adult). Before metamorphosis, embryos and larvae activate their antioxidant pathways and do not show increased oxidative damage. In contrast, post-metamorphic individuals show unnoticeable antioxidant responses, and a marked oxidative damage in lipids. Also, type of pollutant (i.e., organic vs inorganic) promotes contrasting effects across amphibian life stages. Our findings show a divergent evolution of the redox balance in response to pollutants across life transitions of metamorphosing amphibians, likely linked to the characteristics of each life stage. Further comparative mechanistic approaches to wildlife responses to global changes will improve our understanding of these eco-evo-devo processes.

## Introduction

Anthropogenic pollution is considered a major cause of biodiversity loss worldwide (Jaureguiberry et al., 2022). In particular, chemical organic and inorganic pollutants such as fungicides, pesticides, herbicides, or heavy metals are released daily into ecosystems through multiple sources, often resulting in novel and stressful conditions for wildlife (Sigmund et al., 2023). The susceptibility of organisms to these pollutants could change over the course of their lifetime and could be especially important for species with a life cycle that includes abrupt life transitions such as metamorphosis (Lowe et al., 2021). However, no study has systematically evaluated to what extent anthropogenic chemical pollutants impact physiology over the life stages of metamorphosing species. Such assessment would increase our understanding of how chemical pollution impacts on species resilience, which is essential knowledge to develop conservation actions and reduce biodiversity loss.

In low concentrations, reactive oxygen species (ROS) are critical for biological processes since they are involved in immune responses, detoxification and intracellular signalling (Halliwell and Gutteridge, 2015). However, exposure to ROS often causes cell damage, and when the accumulation of ROS overpasses the capacity of antioxidant enzymes to counteract them, a cellular oxidative stress state is induced (Balaban et al., 2005). Oxidative stress can damage essential biomolecules such as lipids, proteins, or DNA, and finally lead to reductions in organismal health and life expectancy (Balaban et al., 2005; Ježek and Hlavatá, 2005). The physiological mechanisms that neutralise ROS (i.e., the redox balance) are thought to have a central systemic role and mediate life-history trade-offs (Costantini, 2019; Costantini et al., 2010; Monaghan et al., 2009; Sohal et al., 2002).

The antioxidant system is highly conserved across taxa and consists of a wide range of enzymatic and non-enzymatic components that work synergistically to control ROS production and achieve redox homeostasis (Pamplona and Costantini, 2011). The first line of defence in response to oxidative damage involves the endogenously produced enzymatic scavengers such as superoxide dismutase, catalase, glutathione peroxidase, or glutathione reductase (Matés and Sánchez-Jiménez, 2000). The second line of defence involves scavenging non-enzymatic antioxidants with a low molecular weight that allows detoxifying ROS located in cellular areas where large enzymes cannot reach (Pamplona and Costantini, 2011). The tripeptide reduced glutathione GSH is the most abundant non-enzymatic molecule in animal cells and can directly scavenge ROS or work in conjunction with antioxidant enzymes (Halliwell and Gutteridge, 2015). The redox balance is therefore set by the action of enzymatic and non-enzymatic antioxidant pathways (Balaban et al., 2005). Finally, some substances are produced under a scenario of oxidative stress as a result of damage to essential biomolecules. Malondialdehyde (an end product of the peroxidation of polyunsaturated fatty acids) is the most common marker of oxidative damage in the lipids of the cell membrane (Mateos and Bravo, 2007). Both the production of ROS and the capacity to neutralise them can be influenced by chemical pollutants via diverse pathways (Benedetti et al., 2015; Valavanidis et al., 2006).

The effect of anthropogenic chemical pollutants on an organism’s redox balance could be linked to an organism’s antioxidant capability as well as its life mode. Both of these likely change across an individual’s ontogeny. This is expected to be particularly relevant across the life cycle of species undergoing metamorphosis, a major life transition undergone by ∼80% of existing animal species (Lowe et al., 2021). Metamorphosing species normally show three remarkably different stages (i.e., embryo, larva [or pupa] and adult) with contrasting phenotypic and physiological characteristics. Among vertebrates, amphibians are an ideal group to study the impact of chemical pollution on the redox balance across contrasting life stages. The life cycle of most amphibians, and particularly of anurans, includes an embryo that hatches into a fish-like larva that abruptly develops a tetrapod juvenile through metamorphosis (Liedtke et al., 2022). Both embryo and larva often have a highly permeable external surface/skin, and their habitat is commonly restricted to the aquatic environment (Alibardi, 2003; Brühl et al., 2011). In contrast, post-metamorphic individuals normally develop a less permeable skin and, although they often rely on waterbodies for breeding, can normally inhabit the terrestrial environment (Alibardi, 2003). Hence, the impact of substances released to water bodies, such as pollutants, is expected to vary across amphibian life stages, with the embryonic and larval stages potentially being the most vulnerable (Brühl et al., 2011). Indeed, amphibians are the most threatened vertebrate group, and chemical pollution is thought to be a relevant force behind their decline (Egea-Serrano et al., 2012). Understanding the extent to which pollutants adversely affect the amphibian redox state at different life stages will add important knowledge for the conservation of these and other metamorphosing species.

During the last three decades, a considerable number of studies have investigated the effect of pollutants on the amphibian redox balance either at their pre- or post-metamorphic stages. Making use of such available information, here we evaluate the impact of anthropogenic chemical pollution (e.g., organic, inorganic pollutants) on different aspects of the redox machinery (enzymatic and non-enzymatic antioxidant responses, and oxidative damage in lipids) in amphibians across their three major life stages (embryo, larva, adult). Specifically, we carry out a systematic literature review and meta-analysis, and assess whether the impact of chemical pollutants on the amphibian redox balance varies across amphibian life stages. We predicted that exposure to pollutants will increase the ROS production in both pre- and post-metamorphic life stages. However, since the external surface of amphibian embryos and larvae is highly permeable and, therefore, can easily absorb water-borne pollutants, we predicted they may have evolved a high antioxidant buffering against pollutants. Consequently, since post-metamorphic amphibians are often less exposed to chemical pollutants due to their high ability to express behavioural plasticity, we predicted a lower antioxidant capacity which would result in high oxidative damage in the presence of chemical pollution. Finally, we expected that the type of pollutant would lead to different consequences on the amphibian redox balance, a process that may be life-stage dependent.

## Results

### Overall effect of pollution on the redox balance of amphibians

An initial meta-analysis (i.e., intercept-only model) including all the oxidative stress parameters and life stages showed that experimental exposure to pollutants increased the levels of redox balance components by 16% compared to control conditions (model intercept [95% confidence interval; ‘95%CI’ hereafter] = 0.150 [0.031, 0.269]; Figure 1a). The total heterogeneity of this model was high (I^2^ = 99.98), with 1.84% and 2.29% of it explained by species and phylogeny respectively, and 25.90% explained by among-study differences.

**Figure 1.**
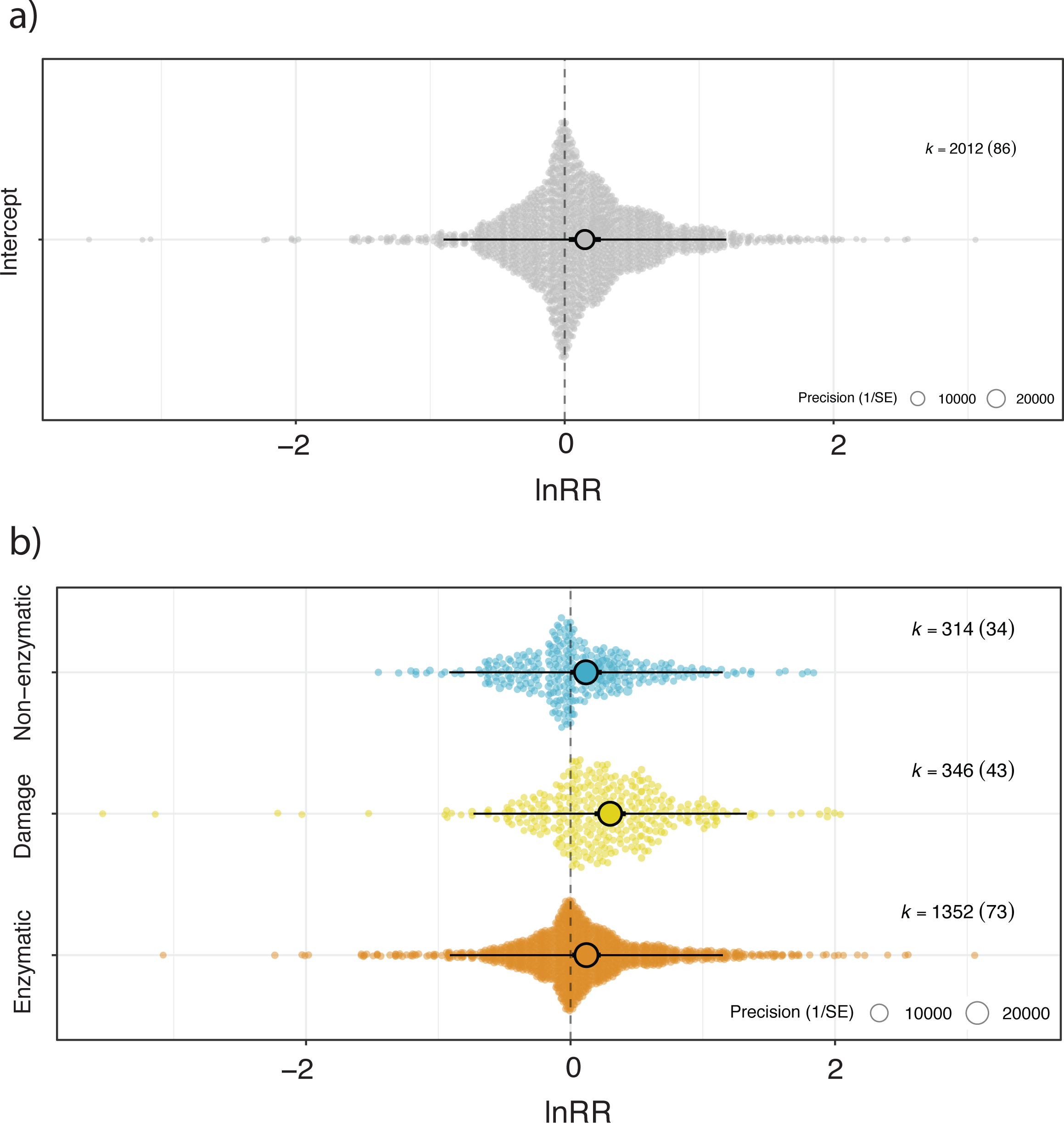
Orchad plots showing the overall effect of pollutant exposure on a) the redox machinery (i.e., pooling all the collected estimates), and b) enzymatic and non-enzymatic parameters, and indicator of oxidative damage in lipids of amphibian species undergoing metamorphosis. Coloured dots show means, thick whisker 95% confidence interval, and thin whisker 95% precision interval. The precision (1/standard error) of each study is represented in the background with scaled grey dots: the bigger the point, the bigger the higher precision. Positive values on the x-axis represent a higher level of a given parameter in response to a pollutant.

Exposure to pollutants increased the levels of all studied redox markers, with 95%CI for model estimates not overlapping zero for enzymatic antioxidants (estimate [95%CI] = 0.120 [0.013, 0.228]; Figure 1) and lipid damage (estimate [95%CI] = 0.299 [0.182, 0.417]; Figure 1), and slightly overlapping zero in the non-enzymatic antioxidants (estimate [95%CI] = 0.117 [-0.002, 0.236]; Figure 1b).

### Effect of chemical pollution across amphibian life stages

Pollutants had a contrasting effect on the redox balance of embryos, larvae, and adults. In embryos, pollutants increased the levels of the non-enzymatic antioxidants (estimate [95%CI] = 0.386 [0.040, 0.733]; Figure 2a) but did not have an effect on the enzymatic antioxidants (estimate [95%CI] = 0.054 [-0.213, 0.320]; Figure 2a) or lipid peroxidation (estimate [95% CI] = 0.221 [-0.205, 0.645]; Figure 2a). Redox marker explained 8.21% of the heterogeneity in redox response to pollutants in embryos (i.e., r^2^_marginal_ = 8.21%). In larvae, pollutants increased to a similar extent the levels of the enzymatic and non-enzymatic antioxidants although not significantly since 95%CI slightly overlapped zero in both cases (estimate ‘enzymatic’ [95%CI] = 0.179 [-0.012, 0.367; Figure 2b] and estimate ‘non-enzymatic’ [95%CI] = 0.200 [-0.013, 0.413]; Figure 2b). In contrast, the effect of pollutants on lipid peroxidation in larvae was very low (estimate [95%CI] = 0.056 [-0.151, 0.264]; Figure 2). Redox marker explained 0.65% of the overall variation in redox response to pollutants in larvae (i.e., r^2^ = 0.65%). In adults, while pollutants had a weak effect both on the enzymatic (estimate [95%CI] = 0.084 [-0.092, 0.261]; Figure 2c) and non-enzymatic antioxidants (estimate [95%CI] = 0.086 [-0.097, 0.268]; Figure 2c), lipid peroxidation levels were substantially increased (estimate [95%CI] = 0.487 [0.304, 0.669]; Figure 2c). Redox marker explained 10.47% of the overall variation in redox response to pollutants in adults (i.e., r^2^_marginal_ = 10.47%).

**Figure 2.**
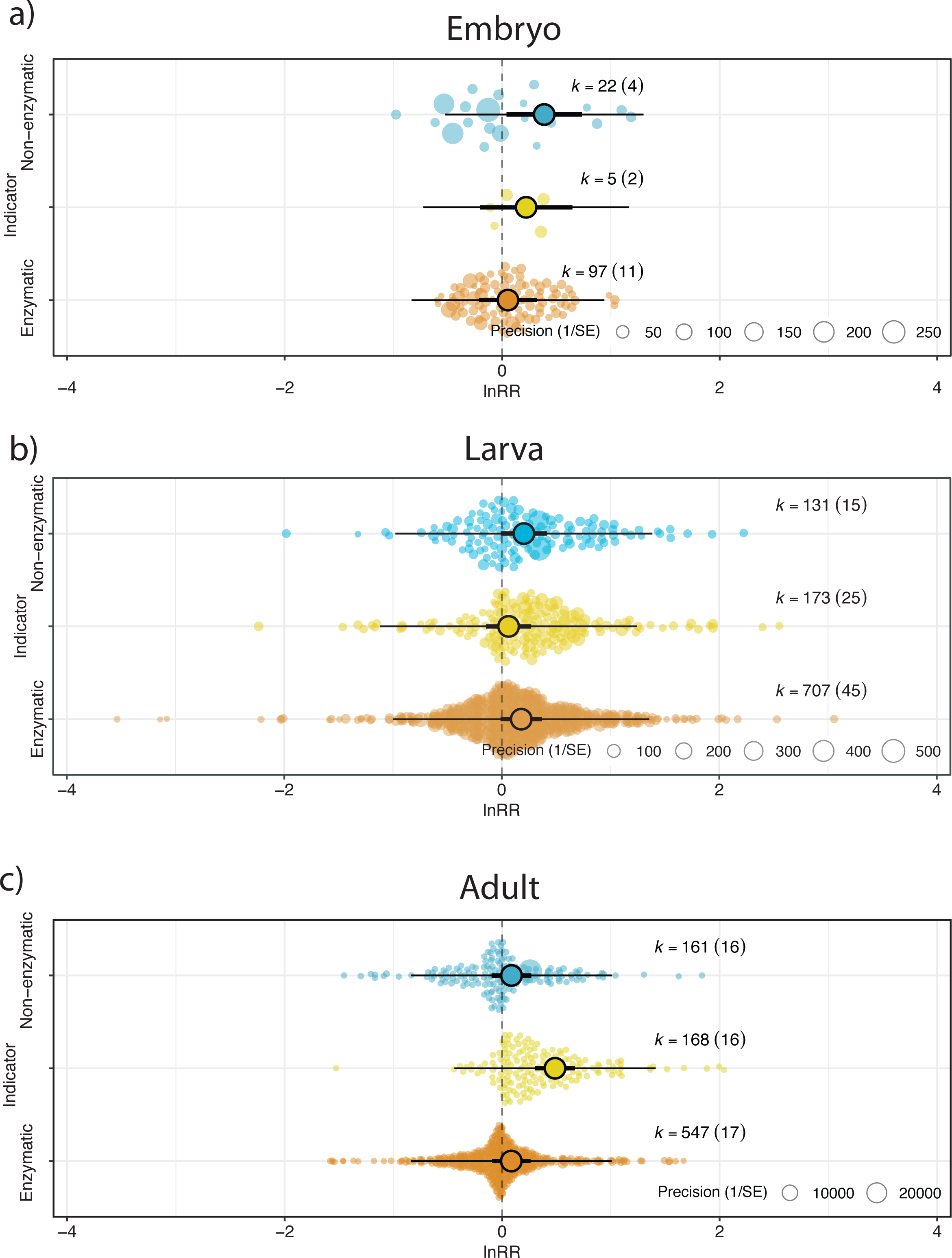
Orchad plots showing the effect of pollutants on the redox machinery (i.e., enzymatic and non-enzymatic components, and indicator of oxidative damage in lipids) of amphibian embryos (a), larvae (b) and adults (c). Coloured dots show means, thick whisker 95% confidence interval, and thin whisker 95% precision interval. The precision (1/standard error) of each study is represented in the background with scaled grey dots: the bigger the point, the bigger the higher precision. Positive values on the x-axis represent a higher level of a given parameter in response to a pollutant.

### Effect of type of pollutants on the amphibian redox state

We first investigated the effect pollutants according to their chemical properties, i.e., organic vs inorganic contaminants. These pollutants only had subtle effects on the redox balance of embryos (Figure 3a). In contrast, organic pollutants increased both the enzymatic and non-enzymatic response in tadpoles with no effect on their lipid peroxidation levels (‘enzymatic’ estimate [95%CI] = 0.236 [-0.006, 0.479], ‘non-enzymatic’ estimate [95%CI] = 0.285 [0.017, 0.553]; Figure 3a) and, in adults, induced lipid peroxidation but no antioxidant responses (‘indicator’ estimate [95%CI] = 0.488 [0.292, 0.683], Figure 3a). The available information for inorganic pollutants is much less; these only had weak effects on the non-enzymatic and enzymatic components of the redox machinery of embryos and adults, respectively (Figure 3b).

**Figure 3.**
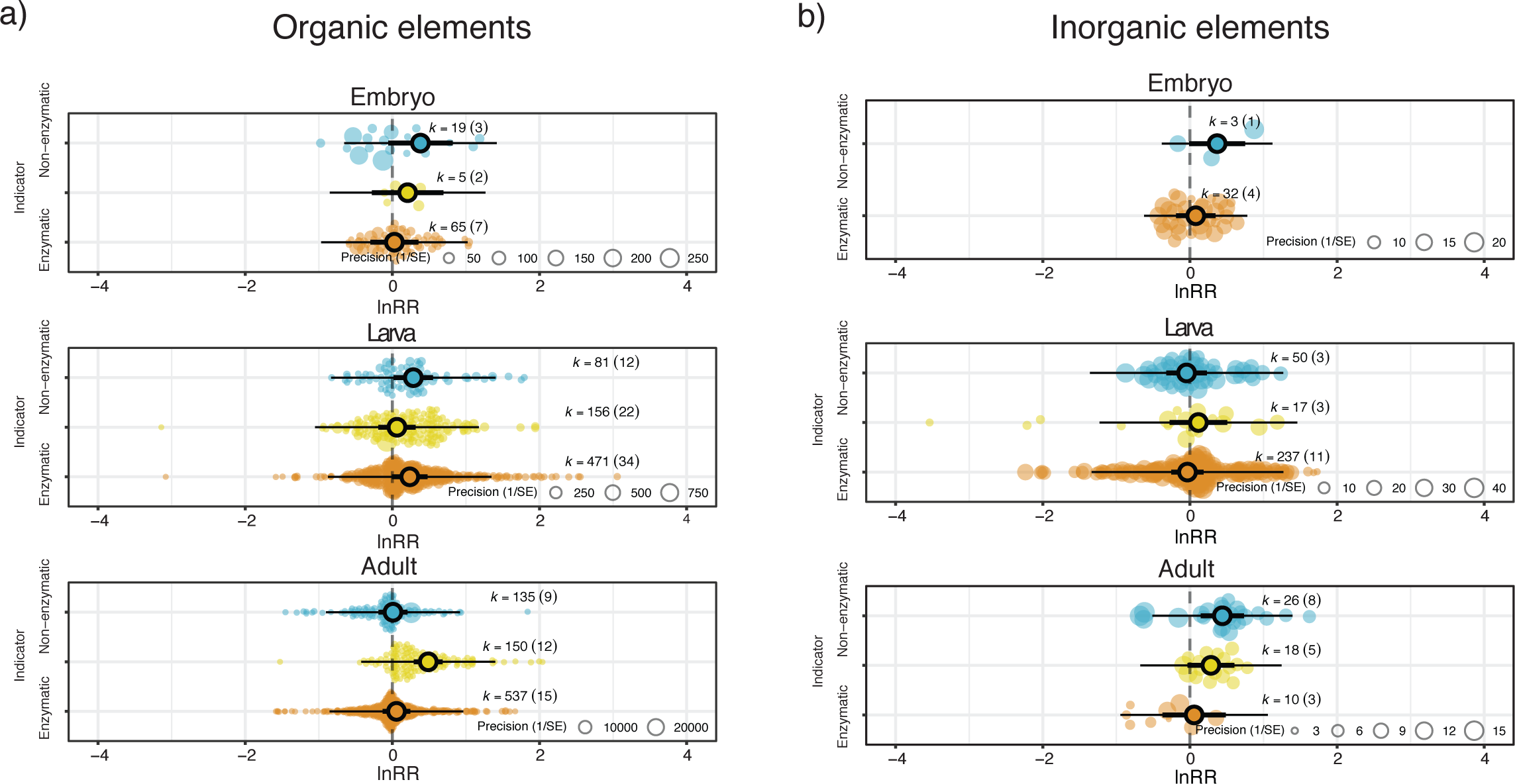
Orchad plots showing the effect of a) organic and b) inorganic pollutants, on the redox machinery (i.e., enzymatic and non-enzymatic components, and indicator of oxidative damage in lipids) of amphibian embryos, larvae, and adults. Coloured dots show means, thick whisker 95% confidence interval, and thin whisker 95% precision interval. The precision (1/standard error) of each study is represented in the background with scaled grey dots: the bigger the point, the bigger the higher precision. Positive values on the x-axis represent a higher level of a given parameter in response to a pollutant.

### Publication bias

We did not detect small-study effects in our dataset (estimate for the square-root of the inverse of the effective sample size [95% CI] = -0.112 [-0.3959, 0.172]), with the overall model intercept after correcting for effective sample size being very similar to the overall model estimate without thais correction (unbiased estimate [95% CI] = 0.215 [0.005, 0.425; overall model intercept presented above = 0.150 [0.031, 0.269]). We did not detect time-lag effects (estimate [95% CI] = 0.004 [-0.010, 0.018]).

## Discussion

Human activities lead to the release of chemical pollutants into ecosystems, which is threatening biodiversity across the globe. Our meta-analysis provides a comprehensive assessment of the impact of chemical pollution on amphibian redox state. Overall, exposure to chemical pollutants increases levels of the key redox biomarkers, indicative of endogenous responses to oxidative stress. However, this effect is life-stage dependent: while embryos and larvae exhibit increased antioxidant responses and experience no oxidative damage to lipids, adults show a lack of antioxidant responses and marked lipid peroxidation. The type of pollutant also has life-stage specific effects on the redox components of amphibians, although the literature is biased towards the effect of specific organic pollutants such as pesticides and herbicides.

The life cycle of most extant animal species includes some form of metamorphosis. This major life transition not only includes conspicuous developmental transformations but also changes in nutrition, niche use and behaviour (Lowe et al., 2021). In metamorphosing amphibians, embryos and larvae have a permeable external surface and are often restricted to water bodies, while post-metamorphic individuals generally have a protective skin and express behavioural plasticity that allows them to explore both the aquatic and terrestrial environments. Our findings shows that the pre-metamorphic antioxidant machinery is upregulated against chemical pollutants, which likely explains the lack of oxidative damage. This pattern contrasts with that observed in post-metamorphic amphibians faced with pollutants, which show negligible antioxidant responses and a marked oxidative damage. Eco-evolutionary differences across amphibian life stages may have driven a divergent ability to cope with variation in environmental pollution. In other words, the likelihood of being exposed to chemical pollutants may have led to a divergent evolution of antioxidant responses across life stages of metamorphosing species (Medina et al., 2007; Saaristo et al., 2018). Oxidative stress is also thought to play a major role in modulating life-history trade-offs, including the balance between important aspects of fitness such as growth and survival (Monaghan et al., 2009). In metamorphosing organisms challenged with pollutants, avoiding an oxidative stress state could allow pre-metamorphic individuals to reach metamorphosis with a body size large enough to reduce post-metamorphic mortality rates (Mccauley et al., 2015; Székely et al., 2020; Zamora-Camacho et al., 2023). Two meta-analyses have suggested that animals show stronger responses to stress at early stages than later in life, which could be a mechanism to avoid developmental impairment and negative carry-over effects (Isaksson, 2010; Messina et al., 2023). However, the evolution of redox responses across an organism’s lifespan may be constrained by its maintaining cost and its relationship with life histories, including phenotypic plasticity (Boonekamp et al., 2018; Burraco et al., 2022). Further empirical and comparative studies will disentangle whether life-stage-dependent redox responses are context- and/or taxa-specific in species undergoing metamorphosis.

Despite chemical pollution commonly leading to negative consequences for wildlife, the mechanistic causes of this process remain unclear. Overall, our study shows that chemical pollutants impact the redox balance of amphibians. These effects vary depending on the type of pollutant and life stage. It should be noted, however, that most of the available data come from studies on the effect of pesticides conducted both in larvae and adults, and herbicides in larvae. With a global production of two million tonnes, chemicals pollutants are ubiquitous in the environment and often enter aquatic ecosystems, posing a major threat to semi-aquatic amphibians (De et al., 2014). The impact of chemical pollutants on amphibian redox balance that we report here might explain, at least partly, the decline of amphibian populations associated with negative chemical pollution (Blaustein et al., 2017). Our study highlights the need for further research on the impact of other chemical pollutants such as metallic elements or other inorganic compounds on the redox balance of amphibians, for which the available information is still scarce. More experimental work to test the effect of these and other pollutants across all life stages will be needed to better understand the real-world impact of chemical pollution on metamorphosing animals. This contrasts with the broad test of those and other pollutants in behavioural studies (Sievers et al., 2019). Our meta-analysis only includes studies using a single pollutant (the most abundant throughout the literature) and experiments combining several pollutants with different characteristics (e.g., chemicals with light or noise pollution) are still scarce, however, they will improve our understanding of the wildlife responses to anthropic disturbed.

### Conclusions

Our meta-analysis shows that the effect of chemical pollution on the redox balance of amphibians varies across the three major life stages of metamorphosing amphibians. While embryos and larvae can induce antioxidant responses to avoid oxidative damage, adults show a lack of antioxidant response but pay a oxidative cost in terms of increased lipid peroxidation. Our study also shows that the type of pollutant can shape the amphibian redox status, which seems to be life-stage dependent. Future studies will provide insights into response to pollutants of different over the developmental trajectory of species undergoing metamorphosis. Experiments designed specifically to examine the link between chemical pollutants, the redox balance and life histories of metamorphosing organisms are needed.

## Methods

### Literature Review

Studies of the effects of anthropogenic chemical pollutants on the redox balance of amphibians were identified via relevant database searches conducted between the 8^th^ of July and the 25^th^ of November 2021. Specifically, using the search string “(‘Oxidative Stress’) AND (‘Amphibians’) AND (‘Pollution’)”, we performed the search on the Web of Science Core Collection, PubMed, EMBASE (Ovid), EBSCOhost, Scopus, and ECOTOX. We read the title and abstract of studies published between 1998 (the earliest year with published data on the meta-analysed topic) and 2021 (when the search was conducted), and we assessed whether studies contained suitable information for our meta-analysis (details below). We identified 865 studies by the database searches above, plus seven additional studies that were identified from the reference list of screened studies (Figure S1). After removing duplicates, 361 studies were screened by reading their title and abstract, and 105 studies were identified as potentially containing suitable information for the meta-analysis. These 105 studies were fully read to assess whether they had information that met the inclusion criteria (see details below). Database searches, study screening and effect size extraction were all performed by one co-author (CM). Most of the data in the papers were presented graphically, and thus numerical data was obtained using the digitalising software WebPlotDigitiser Version 4.4 (Rohatgi, 2013), which has been shown to be a valid and accurate method of data extraction for meta-analyses (Drevon et al., 2017).

### Criteria for inclusion

We were interested in meta-analysing the effects of different pollutants on oxidative stress in amphibians based on controlled laboratory conditions. Therefore, we only included experimental studies that reported: i) mean oxidative stress values, variation (standard deviation or standard error) and sample sizes (i.e., number of individuals) for control (i.e., non-exposed to pollutants) and treatment groups (i.e., exposed to pollutants); ii) one of the following indices markers of the oxidative balance: superoxide dismutase, glutathione peroxidase, catalase, glutathione reductase, glutathione S-transferase (i.e., enzymatic biomarkers), GSH (i.e., non-enzymatic biomarker), or malondialdehyde (i.e., a marker of oxidative damage in lipids); iii) the developmental stage (embryos, larvae or post-metamorphic) in which the effect of pollutants was tested. Additionally, we only included effect sizes from studies that tested one pollutant at a time (i.e., studies not testing the effect of a pollutant in combination with another factor). The full list of pollutants included in this study can be found at the Supplementary Material.

After assessing for inclusion (Figure S1), we extracted 2,012 effect sizes from 86 studies (Acquaroni et al., 2021; Anguiano et al., 2001; Attademo et al., 2016; Awadalla et al., 2019; Barreto et al., 2020; Bhuyan et al., 2020; Burraco and Gomez-Mestre, 2016; Carvalho et al., 2020; Chai et al., 2017; Cheng et al., 2017; Cheron et al., 2022; Costa et al., 2008; Czarniewska et al., 2003; da Silva et al., 2021; David et al., 2012; De Lima Coltro et al., 2017; Dornelles and Oliveira, 2016; Ejilibe et al., 2018; Ezemonye and Tongo, 2010; Falfushynska et al., 2015; Falfushynska et al., 2017; Ferrari et al., 2008; Ferrari et al., 2009; Ferrari et al., 2011; Freitas et al., 2017; Gillardin et al., 2009; Huang et al., 2007; Isnas et al., 2012; Jiang et al., 2019; Jones et al., 2010; Kanter and Celik, 2012; Kostaropoulos et al., 2005; Lajmanovich et al., 2018b; Lajmanovich et al., 2018a; Lajmanovich et al., 2022; Li et al., 2017; Li et al., 2018a; Li et al., 2018b; Liendro et al., 2015; Liu et al., 2006; Liu et al., 2021; Loumbourdis, 2006; Lu et al., 2021; Mardirosian et al., 2015; Marques et al., 2013; Martins et al., 2017; Melvin, 2016; Mussi and Calcaterra, 2010; Naab et al., 2001; Nascimento et al., 2021; Nasia et al., 2018; Özkol et al., 2012; Pal et al., 2018; Papadimitriou and Loumbourdis, 2002; Peltzer et al., 2019; Radovanović et al., 2017; Radovanović et al., 2021; Rosenbaum et al., 2012; Rutkoski et al., 2020; Rutkoski et al., 2021; Saad et al., 2022; Salvaterra et al., 2013; Saria et al., 2014; Shi et al., 2018; Sotomayor et al., 2015; Svartz et al., 2020; Tang et al., 2018; Trachantong et al., 2017; Venturino et al., 2001; Vogiatzis and Loumbourdis, 1998; Wang and Jia, 2009; Wilkens et al., 2019; Wu et al., 2017; Xie et al., 2019; Xu and Huang, 2017; Yin et al., 2014; Yologlu and Ozmen, 2015; Zhang et al., 2012; Zhang et al., 2018a; Zhang et al., 2018b; Zhang et al., 2019b; Zhang et al., 2019a). These 86 studies contained information from 33 amphibian species relatively well distributed across the globe and the amphibian phylogeny (Figure S2 and S3). All these species are anurans with a life cycle including an embryo, larva, and adult (post-metamorphic) stage, and the dataset respectively included 124, 1012, and 876 oxidative stress estimates from these three life stages (see Table S1).

### Meta-analytic effect sizes

We handled the dataset, ran all analyses, and produced visualisations using R (v.4.3.1; R Core Team, 2023). To assess the effects of different pollutants on the oxidative stress of amphibians, we computed the log response ratio (lnRR) (Hedges et al., 1999). We calculated lnRR and its associated sampling variance using the R function ‘escalc’ in the ‘metafor’ R package (v3.8-1; Viechtbauer, 2010). We calculated lnRR so that positive values meant higher values of a given oxidative stress biomarker in the treatment group (i.e., after the exposure to a pollutant) than in the control group (i.e., not exposed to a pollutant), and *vice versa* for negative lnRR values. When a given control group was compared to multiple treatment groups, we divided the sample size of the control group by as many comparisons the control group was used for, and we used this adjusted sample size to calculate lnRR and its sampling variance (98 observations were removed due to a final sample size lower than one). The reported control or treatment standard deviation for 46 observations was zero. These data were retained in the analysis (assigning their standard deviation to 0.01) after checking that their reported SD was correct.

### Meta-analysis

To assess how oxidative stress markers are affected by chemical pollution, we ran a phylogenetic multilevel (intercept-only) meta-analysis and meta-regressions. These models included three random intercept effects: publication identity, phylogeny and species identity, the latter to capture among-species variation not explained by phylogeny. Additionally, an observation identity random term was included to capture variation in effect sizes within studies. For intercept-only models, we estimated total heterogeneity (*I*^2^_total_) (Naganawa and Santos, 2012) and the amount of variation explained by each random term as implemented in the R function ‘i2_ ml’ (‘orchaRd’ R package v2.0; Nakagawa et al., 2021). For meta-regressions, we report on the proportion of variation explained by each moderator as calculated by the R function ‘r2_ ml’ (‘orchaRd’ R package v2.0; Nakagawa et al., 2021).

To understand the overall effect of chemical pollution on redox state, we ran two models. First, we ran an intercept-only model that contained the random effect structure explained above and no moderators. Second, we ran a meta-regression, including the random effect structure presented above and redox biomarker (‘enzymatic antioxidant’, ‘damage’ and ‘non-enzymatic antioxidant’) as a moderator. To understand the effect of chemical pollution across amphibian life stages, we repeated the meta-regression model presented above (i.e., including ‘redox component’ as a moderator) for embryo, larvae and adult life stages separately. To understand the effect of different chemical pollutants across amphibian life stages, we carried out meta-regressions for each type of pollutant (i.e., organic and inorganic) for embryo, larvae and adult life stages separately. These models included the random effect structure presented above. Finally, we also confirmed that the effect of pollutants on the amphibian redox balance was similar across tissues (Figure S4).

### Phylogenies

To control for phylogenetic history, we extracted phylogenetic trees from Open Tree of Life (Hinchliff et al., 2015; Rees and Cranston, 2017), accessed via the R package ‘rotl’ (v3.0.14; Michonneau et al., 2016). Tree branch length was calculated following Grafen (1989) and we generated a phylogenetic correlation matrix that was included in all our models. We assessed the phylogenetic importance in our meta-models calculating the proportion of variation in lnRR explained by the phylogeny (*I*^2^_phylogeny_) (Cinar et al., 2022).

### Publication bias

We tested small-study effects and time-lag effects following Nakagawa et al. (2022) by running two additional multilevel meta-analytic models of lnRR. Each of these models included, as a single moderator, either the square-root of the inverse of the effective sample size or the mean-centred year of study publication (Ioannidis and Trikalinos, 2005; Nakagawa et al., 2022).

## Acknowledgements

PB was supported by Marie Sklodowoska-Curie Individual Fellowship 797879 METAGE and by Juan de la Cierva Incorporación IJC2020-044682-I (Spanish Ministry of Science and Innovation). PM was supported by ERC Advanced Grant 101020037 under the European Union’s Horizon 2020 research and innovation program.

## Statement of authorship

PB and PM conceived the study; CM performed data extraction, CM and PC-L carried out all the statistical analysis; CM and PB wrote the first draft of the manuscript and all authors contributed substantially to the final version of the manuscript.

## Data statement

All R scripts and datasets needed to reproduce the analyses presented in this paper are now available at: https://github.com/PabloCapilla/meta-analysis_pollution. Should the manuscript be accepted, a DOI to to a public repository will be provided.

## Notes

### Competing Interest Statement

The authors have declared no competing interest.

