## Supplementary material for "The impact of chemical pollution across major life transitions: a meta-analysis on oxidative stress in amphibians": Suplementary Material

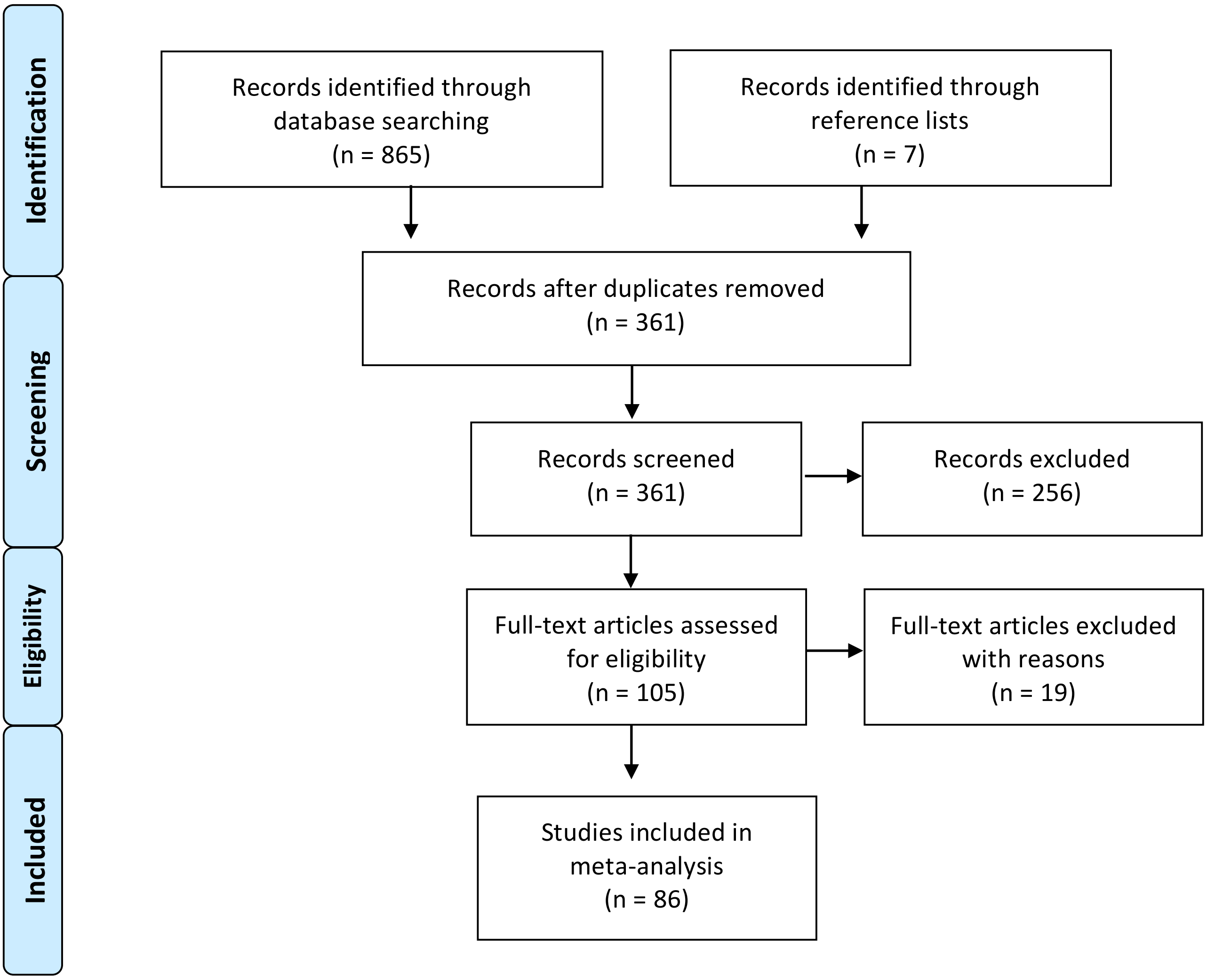


**Figure S1.** PRISMA Flow Diagram outlining literature review process.

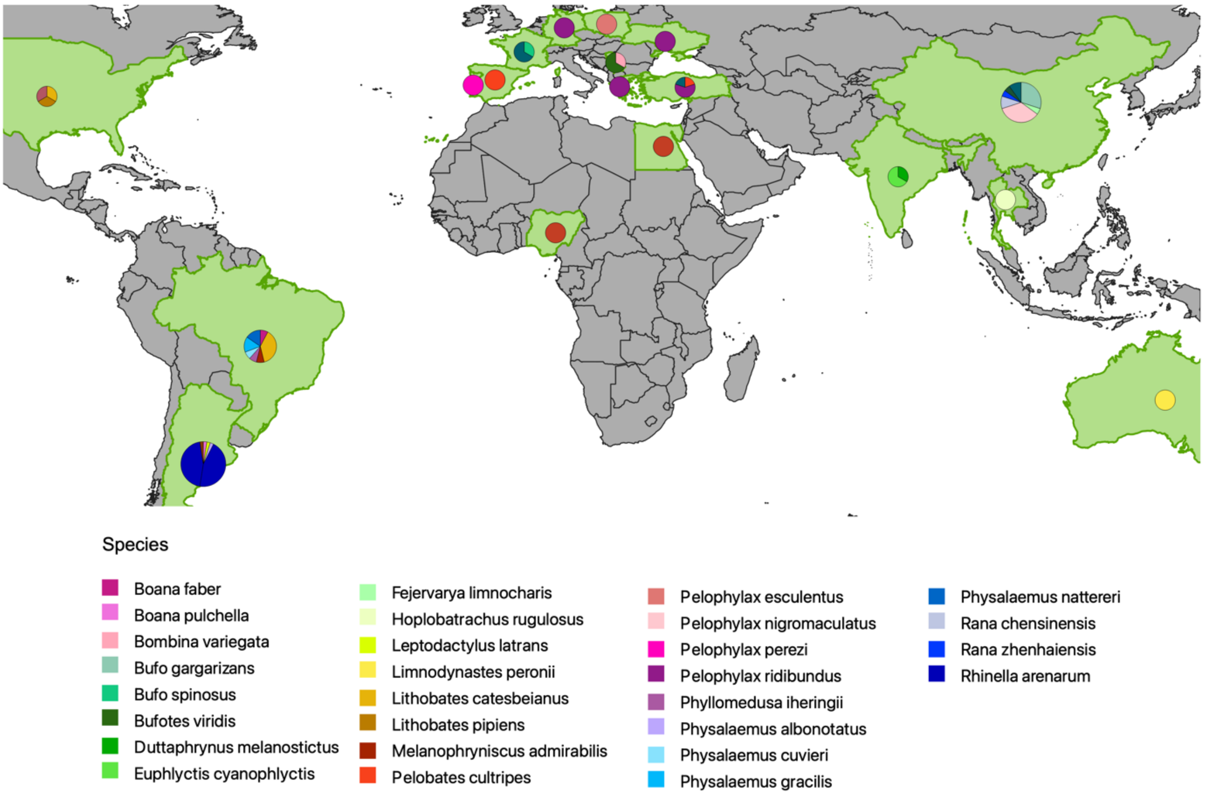


**Figure S2.** Map showing the countries where the studies included in the meta-analysis were carried out (in green), and the species used in each case (circules, see color legend).

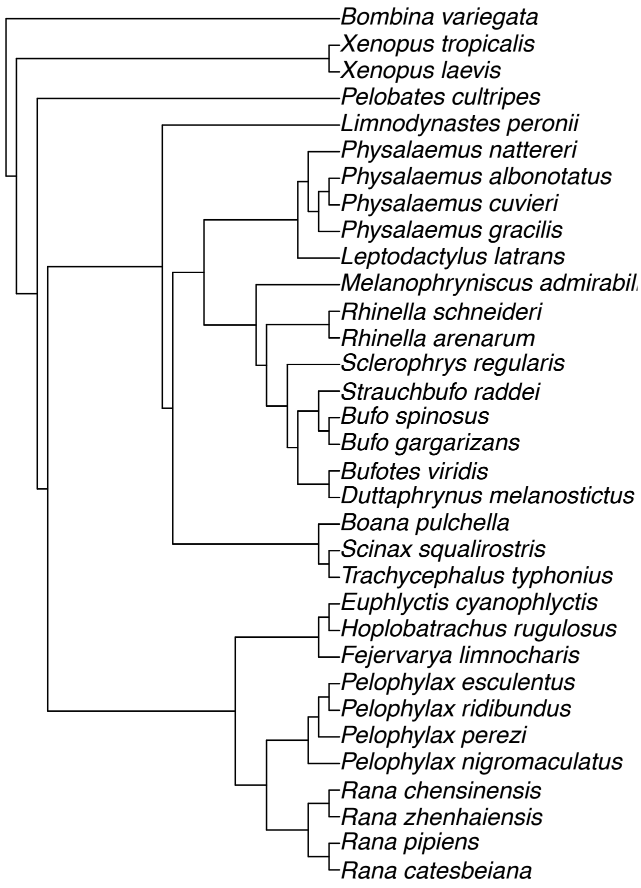


**Figure S3.** Phylogeny of the species included in the meta-analysis.

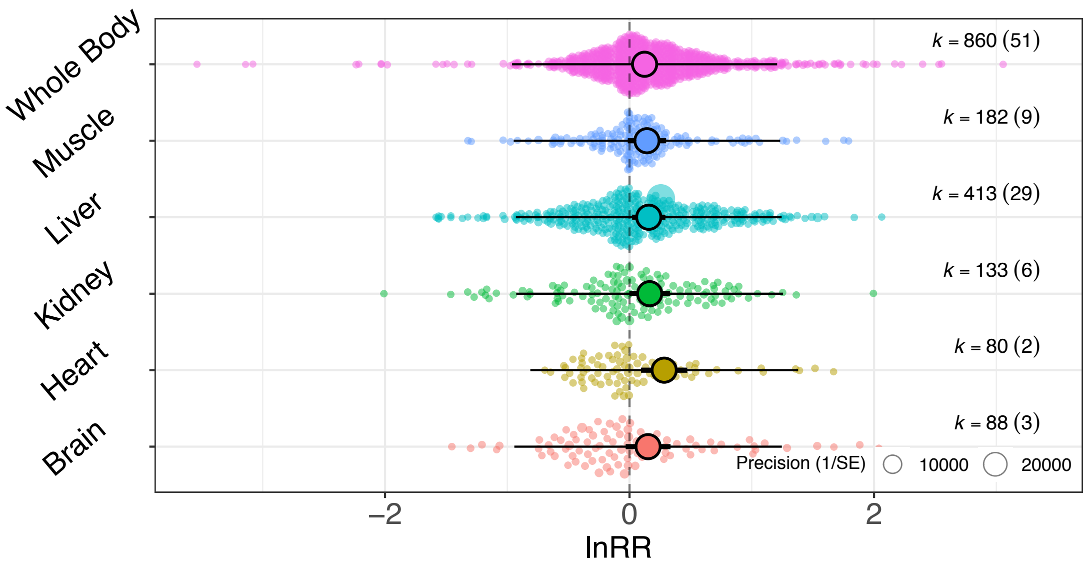


**Figure S4.** A meta-regression model assessing redox responses across the main tissues included in the meta-analysis dataset (pooling data from different life stages and redox components) showed a similar change among them (estimate [95%CI]: whole body = 0.123 [0.014, 0.232]), liver = 0.157 [0.021, 0.294], muscle = 0.143 [-0.015, 0.300], kidney = 0.165 [-0.002, 0.330], brain = 0.152 [-0.032, 0.335], and heart = 0.284 [0.094, 0.473]). Tissue type only explained 0.39% of the overall variation in redox response to pollutants.

| | | Organic compound | Inorganic compound |
|---|---|---|---|
| **Embryo** | **Enzymatic** | 65 (7) | 32 (4) |
| | **Non-enzymatic** | 19 (9) | 3(1) |
| | **Lipid damage** | 5 (2) | 0 (0) |
| **Tadpole** | **Enzymatic** | 471 (34) | 237 (11) |
| | **Non-enzymatic** | 81 (12) | 50 (3) |
| | **Lipid damage** | 156 (22) | 17 (3) |
| **Adult** | **Enzymatic** | 537 (15) | 10 (3) |
| | **Non-enzymatic** | 135 (9) | 26 (8) |
| | **Lipid damage** | 150 (12) | 18 (5) |

**Table S1.** Number of estimates and studies (in brackets) across developmental stages, redox balance traits and type of pollutants.

**List of pollutants included in this study.**

Pesticides (Aminomethylphosphonic acid, azoxystrobin, carbofuran, metaldehyde), fungicides (myclobutanil, cyproconazole, difenoconazole, epoxiconazole, metalaxyl, tebuconazole, triadimefon, triadimenol), herbicides (glyphosate, 2,4-dichlorophenoxyacetic acid, acetochlor, atrazine, butaforce, Roundup Ultra-Max, C-K Yuyos FAV, clomazone, glifoglex, glufosinate-ammonium, Infosato, metamifop, paraquat, quinclorac, sulfentrazone), Insecticides (lambda-cyhalothrin, 3,3′,4,4′,5-pentachlorobiphenyl, acephate, azinphos carbaryl, azinphos methyl, bacillus thuringiensis israelensis, carbaryl, chlorpyrifos, cypermethrin, deltamethrin, diazinon, dieldrin, diflubenzuron, dimethoate, endosulfan, fenthion, fipronil, lindane, malathion, methamidophos, methomyl, metoxychlor, neonicotinoid (acetamiprid), neonicotinoid (Thiamethoxam), octylphenol, omethoate, parathion, permethrine, bifenthrin, spirotetramat, trichlorfon), metallic elements (aluminium, arsenic, barium, beryllium, cadmium, calcium, chromium, cobalt, copper, gallium, indium, iron, lead, manganese, mercury, nickel, potassium, rubidium, strontium, uranium, zinc), pharmaceuticals (dexamethasone, diclofenac, naproxen, atenolol, gemfibrozil, nifedipine, polyethylene glycol), wastewater contaminants (4-Methylbenzylidene camphor, astrazon dyes, cibacron dyes, phenanthrene, polyfluoroalkyl chemicals, remazol dyes, sodium fluoride, triclosan, zinc oxide), nanoparticles (Ni/<gamma>-Al2O3, <gamma>-Al2O3, NiO/<gamma>-Al2O3, silicon dioxide, titanium silicate), and Multiwalled Carbon NanoTubes.
